# ColiFormer: A Transformer-Based Codon Optimization Model Balancing Multiple Objectives for Enhanced *E. coli* Gene Expression

**DOI:** 10.1101/2025.11.26.690826

**Authors:** Saketh Baddam, Omar Emam, Abdelrahman Elfikky, Francesco Cavarretta, George Luka, Ibrahim Farag, Yasser Sanad

## Abstract

Codon optimization has become a key strategy to improving heterologous gene expression in *Escherichia coli*. However, many existing methods focus primarily on maximizing the codon adaptation index (CAI) while overlooking the importance of codon harmonization, which is crucial for maintaining proper protein characteristics.

In this study, we present ColiFormer, a transformer-based codon optimization framework that was fine-tuned on 3,676 high-expression *E. coli* genes curated from the NCBI database. Built upon the CodonTransformer BigBird architecture, ColiFormer employs self-attention mechanisms and a mathematical optimization method (the augmented Lagrangian approach) to balance multiple biological objectives simultaneously, including CAI, GC content, tRNA adaptation index (tAI), RNA stability, and minimization of negative cis-regulatory elements.

Performance was evaluated on 37,053 native *E. coli* genes and 80 recombinant protein targets commonly used in industrial studies, and the results were compared with six established codon optimization approaches. Across all datasets, ColiFormer demonstrated significant improvements in CAI and tAI, maintained GC content within optimal ranges, and reduced the incidence of negative cis-regulatory elements, all with lower runtime costs than most alternative methods.

These results, based on in silico metrics, indicate that ColiFormer consistently enhances recombinant protein expression and achieves superior performance across multiple benchmarks compared to other established techniques. The tool is available as an open-source software package, along with the benchmark sequences dataset used in this study.

## 1. Introduction

Since the seminal breakthroughs of Chang and Cohen (1974) and Morrow et al. (1974), heterologous gene expression has become a cornerstone of modern biotechnology [1,2]. The expression of recombinant proteins in a heterologous host has a wide range of applications, including therapeutic protein production, vaccine manufacturing, diagnostic assays in biomedical fields, bioremediation in industrial fields, and food processing [3, 4, 5, 6]. In particular, *E. coli* is a widely used host factory for protein expression, demonstrated by its use in producing vaccines (e.g., the malaria vaccine FALVAC-1) and other recombinant proteins [7].

The recombinant protein therapeutics market reached $3.01 billion in 2024 and is projected to grow at a CAGR of 10.2% to $5.58 billion by 2030 [8]. However, codon bias is reducing yield, affecting protein quality, and reducing the expression yield of several heterologous proteins–often falling below 0.1 mg/ 100 mL of culture [9]. The redundancy of the genetic code (61 sense codons for 20 amino acids) allows for multiple codons–synonymous codons–to encode the same protein; however, organisms exhibit codon bias by preferentially using certain synonymous codons over others. This bias arises due to differences in tRNA (Transfer Ribonucleic Acid) abundance and the efficiency of the translation machinery [10].

Rare codons, despite their potential to slow translation, play a crucial role in enhancing protein folding efficiency by introducing translational pauses that allow nascent polypeptides to correctly adopt their functional conformations. These pauses appear because the ribosome has to wait longer for low-abundance tRNAs, which briefly slows elongation and gives nearby regions of the polypeptide a realistic chance to settle into the structures they form early on. Conversely, abundant codons promote faster and more accurate translation rates, improving overall protein yield but sometimes at the cost of misfolding or aggregation due to insufficient translational folding time [11]. When decoding is rapid, the chain can move forward before those early structures form fully, which increases the load on chaperone systems.

Codon harmonization solutions attempt to resolve this dilemma by recreating the native gene’s codon usage patterns in the heterologous host. Rather than just increasing the use of abundant codons, harmonization balances synonymous codon selection to improve translational kinetics, ensuring appropriate ribosomal pausing when required, allowing accurate folding and functional protein expression [12]. Keeping certain slow-decoding sites helps maintain the native timing of translation while still improving expression in the rest of the sequence. Aligning with codon bias, the selection of synonymous codons that are more commonly used in the host genome can significantly enhance protein expression, with improvements ranging from 2- to 15-fold in *E. coli* [13].

However, despite their potential to slow translation, rare codons can enhance the folding efficiency of the protein, allowing proper protein expression and functional conformation [14]. Other studies observe that codon bias–while it increases expression–introduces unintended issues: cellular toxicity, tRNA imbalance, and metabolic stress, which ultimately hinder protein production, solubility, and stability [11, 15]. In heterologous contexts, this mismatch between gene codon usage and host tRNA pools directly affects protein yield and folding fidelity [13]. This trade-off between rare and abundant codons makes it necessary to strategically use codon optimization for recombinant genes.

Codon optimization seeks to balance codon usage preferences with protein production goals to maximize yield without compromising the quality and functionality of the expressed protein. It involves strategically modifying codons to align with the host organism’s preferences while preserving the original amino acid sequence [16, 17]. Numerous biotechnology companies have developed extensive codon optimization algorithms to address these challenges. However, these algorithms have a major limitation: they focus predominantly on maximizing protein production and CAI (Codon Adaptation Index), neglecting the biological context and characteristics of the gene [12]. This narrow focus on protein yield frequently results in failures in heterologous expression, leading to costly expression failures and reduced yield. Furthermore, the lack of multiple objectives balancing and inability to capture long-range dependencies within sequences cause imbalances in tRNA pools, leading to cellular toxicity and corrupted protein folding and function [10, 14]. This neglect of the complex effects of codon bias within the host cell has prompted the rise of machine learning and deep learning methods, which better capture the complex relationships underlying codon usage and expression outcomes.

As an alternative to existing approaches, we introduce a new method, ColiFormer, an *E. coli*-specific transformer-based codon optimizer, built upon the BigBird CodonTransformer architecture, designed to balance codon adaptation and biological constraints to improve heterologous protein expression in *E. coli.* [18]. Fine-tuned on 3,676 high-CAI *E. coli* genes curated from NCBI (National Center for Biotechnology Information), ColiFormer leverages the power of self-attention and fine-tuning to adapt from general sequence understanding to *E. coli*-specific codon usage, accounting for the complexity of codon bias within the host cell. It enables simultaneous control of multiple objectives (e.g., CAI, GC content, tAI) through Lagrangian optimization, outperforming current AI tools such as NCRF (Neural Codon Random Forest), CodonTransformer, and CodonGPT –as demonstrated in later sections [12, 18, 19]. ColiFormer addresses the critical trade-offs between codon optimization and codon harmonization and offers an outperforming, AI-driven solution for industrial gene synthesis and therapeutic protein production.

To evaluate ColiFormer’s performance, we benchmarked it on 80 industry-relevant genes derived from past recombinant protein expression studies, alongside 37,053 *E. coli* genes. ColiFormer’s optimized outputs were compared against six established methods: the original (unoptimized) sequence, IDT, Extended Random Choice (ERC), CodonGPT, NCRF, and CodonTransformer. Optimization performance was quantified using multiple in silico metrics, including CAI, GC content (Guanine-Cytosine content), negative cis-regulatory elements, RNA stability, tAI (tRNA Adaptation Index), success rate, and algorithm run-time.

## 2. Materials and Methods

### 2.1. Database and Benchmark Set

With 349,154 *E. coli* genomes, each one containing ∼ 4400 genes, the National Center for Biotechnology Information’s (NCBI) GenBank database was used to collect the *E. coli* gene sequences [20]. Then, CD-HIT-EST [21] was used to cluster highly similar sequences that shared greater than 90% sequence identity, thereby reducing redundancy and resulting in 41435 unique gene sequences listed in Supplemental File 1. To ensure the biological validity of the gene dataset, further cleaning steps–removing sequences with internal stop codons and confirming the presence of valid start and stop codons–were performed, as shown in Figure 1. To focus on highly expressed genes, the remaining sequences were ranked based on their CAI, calculated using a standard reference set of highly expressed *E. coli* genes. Listed in Supplemental File 2, the top 3,676 high-CAI sequences were selected as the final training database, reflecting the typical codon-usage patterns and native characteristics of highly expressed *E. coli* genes. The remaining 37,053 gene sequences were reserved to evaluate ColiFormer’s ability to optimize native *E. coli* sequences.

**Figure 1.**
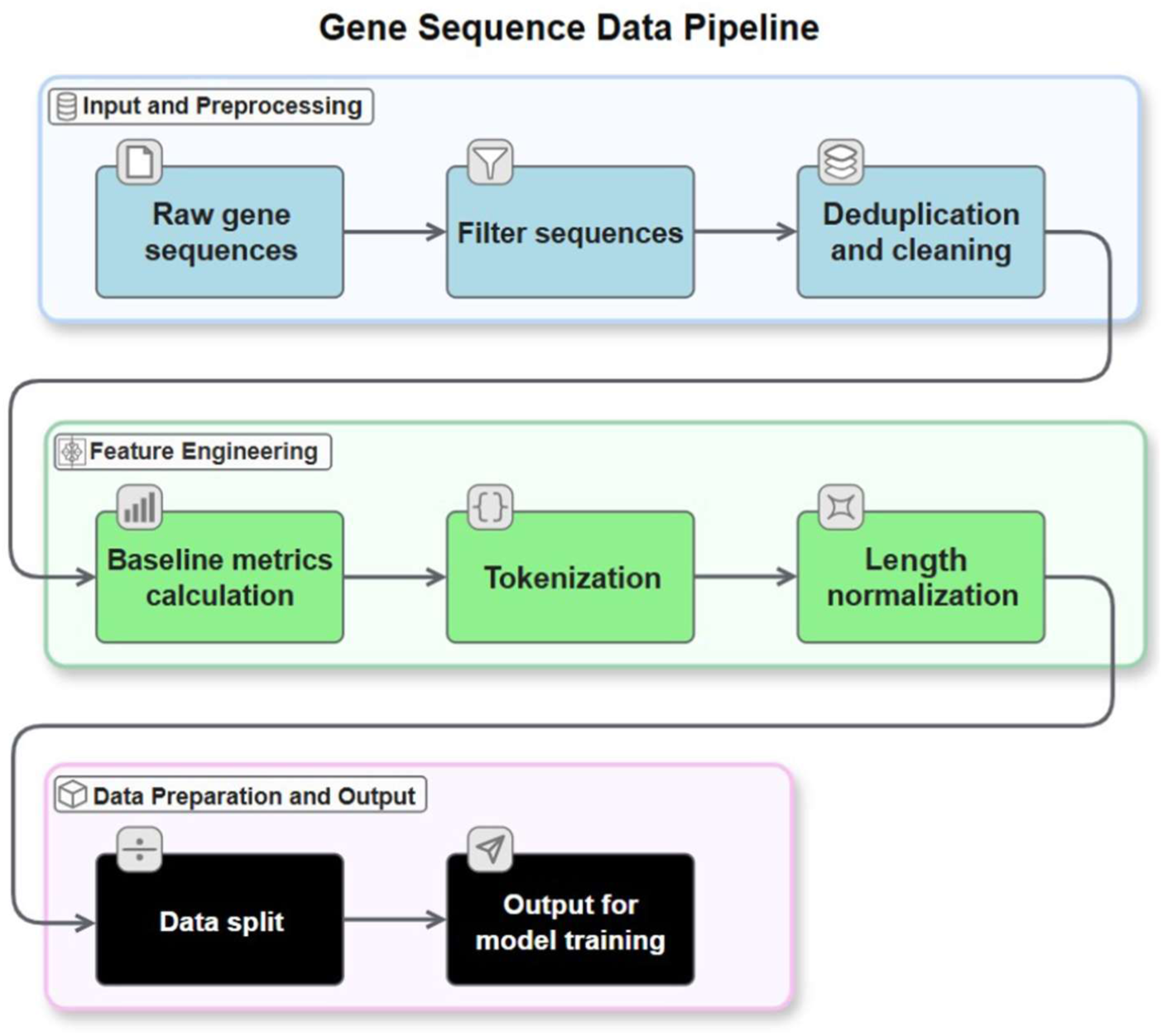
This diagram illustrates the complete data preprocessing workflow for ColiFormer. Starting with raw *E. coli* gene sequences, the pipeline includes filtering incomplete sequences, duplication to remove redundancies, and cleaning steps. Feature engineering follows, where baseline metrics like CAI and GC content are calculated, sequences are tokenized into codon units, and length normalization is applied. The processed data is then split into training and validation sets, ready for input into the model training process

As a benchmark set, 80 DNA sequences were compiled from published datasets and studies focused on heterologous protein production in *E. coli*. These genes cover a broad range of functional categories to ensure the fine-tuned model’s generalizability. This benchmark set serves as a validation for the tool’s effectiveness in optimizing well-established recombinant targets. These 80 DNA sequences, along with their sources, are provided in Supplemental File 3.

### 2.2. CodonTransformer Model Architecture

CodonTransformer is a multispecies deep learning model initially trained on over 1 million DNA-protein pairs from 164 organisms across all domains of life [18]. It employs a Transformer architecture with advanced context-awareness, using a sequence representation method called STREAM (Shared Token Representation and Encoding with Aligned Multi-masking). STREAM encodes both codon and amino acid identity per token, enabling the model to capture organism-specific codon preferences while preserving protein-level constraints. CodonTransformer study shows that this representation generalizes well to organisms not included in the training dataset, due to the model’s reliance on the codon-usage patterns learned across diverse species.

The model’s base architecture includes 89.6 million parameters built upon the BigBird Transformer variant, comprising 12 hidden layers with 768 hidden dimensions, 12 attention heads, and a block sparse attention mechanism with a block size of 64. BigBird was selected for its efficiency in modeling long-range dependencies in long, extensive genetic sequences, which is essential for capturing codon context [22]. This architecture enables efficient processing of sequences up to 2,048 tokens while maintaining computational tractability through sparse attention patterns. The model’s 2,048-token maximum input length is not a constraint here, since *E. coli* genes are shorter than this threshold [23]. Figure 2 illustrates the ColiFormer architecture built upon CodonTransformer, highlighting token embeddings, frozen and trainable transformer blocks, and multi-objective sequence evaluation and generation methods.

**Figure 2.**
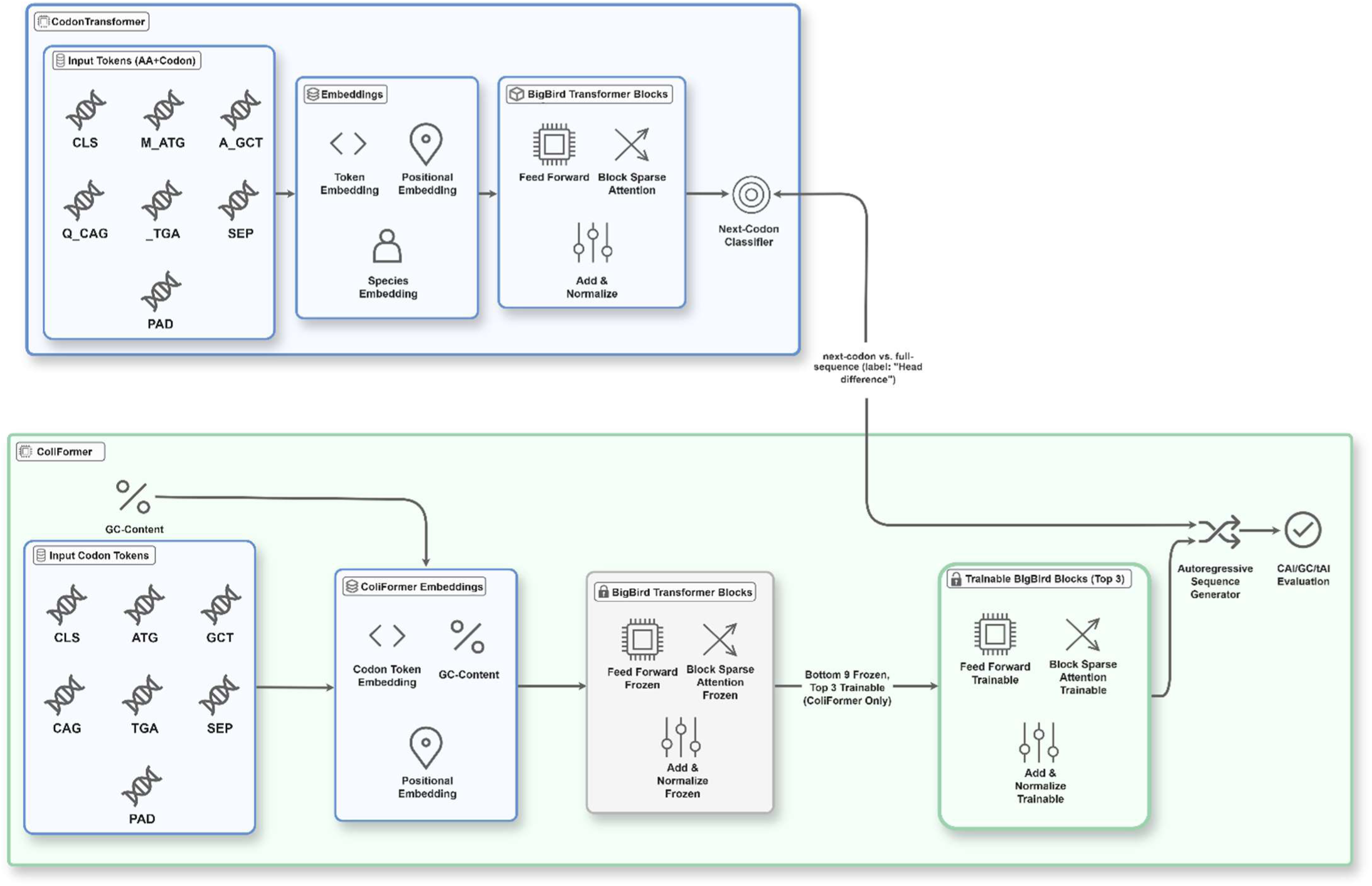
This diagram depicts the ColiFormer system built upon the CodonTransformer architecture. Input codon tokens are embedded with position and GC-content information before passing through a series of transformer blocks. The model distinguishes between frozen and trainable blocks, allowing pretrained knowledge with targeted fine-tuning. The output sequences undergo autoregressive generation and multi-objective evaluation, facilitating precise codon optimization tailored for *E. coli* gene expression.

A key innovation of CodonTransformer is its dual representation strategy, where each token simultaneously encodes amino acid identity and codon information. Its vocabulary consists of 90 distinct tokens, including 64 amino acid-codon combinations, 20 masked amino acid tokens (a_unk through y_unk), 3 stop tokens, and 5 special tokens ([UNK], [CLS], [SEP], [PAD], [MASK]) for sequence processing. This representation enables the model to simultaneously learn amino acid sequence constraints and organism-specific codon usage patterns through masked language modeling objectives.

### 2.3. Dataset Preparation and Curation

For *E. coli*-specific optimization, we curated a high-quality dataset from multiple sources, selecting 3,676 genes with elevated CAI values for model fine-tuning. This dataset primarily includes *E. coli* genes with documented high expression characteristics, validated against an established reference database of high-expression genes. Each sequence underwent rigorous validation to ensure structural integrity, including verification of proper start codons (ATG, TTG, CTG, or GTG; present in their natural frequencies), appropriate stop codon termination (TAA, TAG, or TGA), absence of internal stop codons, sequence length divisibility by three, and exclusive composition of standard nucleotides (A, T, G, C). All sequences passed these validation checks, with none excluded.

The validated sequences were processed into the CodonTransformer input format, where protein sequences were represented as masked token sequences [24]. For instance, a protein sequence “MALW” would be initially represented as “M_UNK A_UNK L_UNK W_UNK”, with UNK indicating unknown codon assignments. During training, the model learns to predict the optimal codons at each position based on surrounding sequence context and *E. coli* usage patterns. During inference, these masked tokens are resolved into context-optimized codons using maximum probability decoding or beam search methods. All sequences were tagged with organism ID 51, corresponding to “*Escherichia coli* general” in the CodonTransformer organism registry. The data preprocessing pipeline, including raw sequence filtering, deduplication, feature extraction, and preparation of inputs for model training, is summarized in Figure 1.

### 2.4. Fine-tuning Methodology with Enhanced GC Control

A critical innovation in ColiFormer is the implementation of an enhanced Augmented-Lagrangian Method to achieve precise control of GC content during fine-tuning [25]. In *E. coli*, maintaining an optimal GC content of around 52% is crucial, as deviations can negatively impact gene expression efficiency and cellular fitness, as shown in Figure 2. Deviation from optimal GC content in *E. coli* may disrupt mRNA secondary structure, impair ribosomal binding, and reduce transcript stability. Traditional codon optimization approaches often struggle to maintain appropriate GC balance while optimizing for other parameters, resulting in sequences with suboptimal expression.

To address this, we formulated the optimization problem as a constrained learning task where the model minimizes masked language modeling loss while enforcing GC content constraints. The total loss function is formulated as:

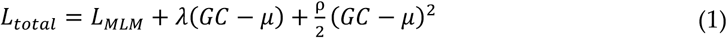

Here, 𝐿_*MLM*_ represents the (masked language modeling) loss; when the constraint terms in the objective function are weak, this encourages a rise in the CAI, but the GC content may drift from the target. λ represents the Lagrangian multiplier; it corrects directional bias by shifting probabilities toward lower-GC codons when the GC content exceeds the target, or toward higher-GC codons when it is below the target. GC is the predicted GC content calculated through differentiable operations, and μ equals 0.52 as the target GC content for *E. coli*. ρ is the adaptive penalty coefficient that controls the strictness of GC adherence.

The MLM loss is defined as follows:

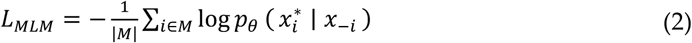

Here, 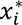 is the gold codon at position 𝑖; this is the true codon to be predicted at the masked location. ∣ 𝑀 ∣ is the total number of masked positions in the input sequence. 𝑥_−*i*_ is the sequence context excluding position 𝑖; all codons in the input except for the masked codon at 𝑖. 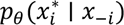 is the probability, according to the model’s parameters 𝜃, of the gold codon 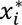 occurring at position 𝑖 given the masked input context 𝑥_−*i*_.

The GC content calculation leverages a differentiable lookup table mapping each token in the vocabulary to its corresponding GC fraction. For any predicted probability distribution over tokens, the expected GC content is computed as the weighted sum of individual token GC values. This differentiable formulation enables gradient-based optimization while maintaining biological constraints. To ensure robust and stable convergence, ColiFormer implements a self-tuning mechanism for the penalty coefficient 𝜌, which adaptively adjusts the strictness of GC content constraints during training. The deviation from the target GC content is defined as Δ*_k_* = 𝐺𝐶*_sm_* − 𝜇, where 𝐺𝐶*_sm_* is the smoothed, differentiable GC estimate and 𝜇 is the desired GC value. The violation at the step 𝑘 is quantified as 𝑣*_k_* =∣ Δ*_k_* ∣. After an initial curriculum warm-up phase (first 3 epochs), the loss incorporates augmented-Lagrangian terms, and both the Lagrange multiplier (𝜆) and penalty coefficient (𝜌) undergo scheduled updates every 20 steps. Specifically, at each scheduled update, the dual variable is updated via:

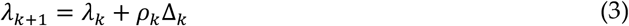

Meanwhile, the penalty coefficient is adaptively increased when constraint violations persist, using the following rule:

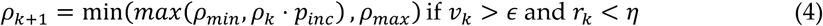

Where 𝜖 is the violation tolerance, 𝑝*_ine_* is the penalty update factor, 𝑟*_k_* is the relative improvement threshold, and 𝜌*_min_*, 𝜌*_max_* set allowed bounds for 𝜌. Otherwise, 𝜌*_k_*_+1_ = 𝜌*_k_*. All parameters and violations are logged at each step.

This adaptive update mechanism tightens the GC constraint only when the model’s progress toward the target GC is inadequate, stabilizing 𝜌 when the target is closely matched. As a result, ColiFormer maintains precise control over sequence composition, allowing robust optimization even when codon context and GC control are in conflict.

### 2.5. Training Configuration and Implementation

Training was conducted on NVIDIA GPU infrastructure with 16GB VRAM (Video Random Access Memory (GPU memory)) using mixed precision computation to maximize efficiency [26]. The model was fine-tuned for 15 epochs with a batch size of 6 sequences, employing the AdamW optimizer with a learning rate of 5 × 10⁻⁵. We implemented a cosine annealing schedule with warm restarts, where the learning rate periodically returns to its initial value before decaying again, promoting exploration of the parameter space and preventing premature convergence to suboptimal solutions [27].

The number of fine-tuning epochs (15) was selected based on early convergence of (i) the validation masked language modeling loss and (ii) the GC-content constraint violation metric (V = ∣GC − μ∣). During training, both metrics were monitored at each epoch. We observed that the loss and GC violation typically plateaued, showing less than 0.5% improvement by epoch 13, indicating satisfactory convergence without overfitting. To ensure complete convergence across all runs, we set the total number of epochs to 15.

A curriculum learning strategy was employed where GC constraints were enforced only after a 3-epoch warm-up period. This approach allows the model to first adapt to *E. coli*-specific codon usage patterns before introducing the additional constraint of GC content optimization. During the warm-up phase, the model focuses solely on minimizing masked language modeling loss, learning the intrinsic patterns of high-expression *E. coli* genes. After the curriculum period, the augmented Lagrangian terms are activated, guiding ColiFormer toward sequences that satisfy both expression optimization and GC content requirements. Validation loss plateaued at epoch 13, indicating convergence without overfitting.

The training process incorporated extensive monitoring of both standard metrics and constraint satisfaction. The Lagrangian multiplier λ was updated every 20 optimization steps based on the current constraint violation, while the penalty coefficient ρ underwent adaptive adjustment based on the relative improvement in constraint satisfaction. This dual update mechanism ensures robust convergence even for difficult optimization landscapes where codon preferences and GC constraints may conflict.

### 2.6. Sequence Generation and Inference Strategies

The fine-tuned model supports multiple sequence generation strategies to accommodate different objectives and constraints. For standard optimization tasks, deterministic decoding selects the highest probability codon at each position, producing consistent and reproducible sequences. For applications requiring sequence diversity, stochastic sampling with temperature scaling enables the generation of multiple distinct sequences while maintaining optimization quality [28]. A key feature is the implementation of a constrained beam search algorithm that maintains multiple candidate sequences simultaneously while enforcing strict GC content bounds throughout the generation process. This approach prevents the overuse of high-frequency codons and ensures the biological viability of optimized sequences. Compared to standard beam search, this constrained version maintains similar computational cost while ensuring that candidates stay within biological limits.

The constrained beam search algorithm incorporates position-aware GC tracking, where each partial sequence maintains its cumulative GC count and projected final GC content. Candidate expansions that would violate the specified GC bounds are pruned from the search space, ensuring all generated sequences satisfy the biological constraints. In practice, increasing beam size expands the search space and can improve sequence quality, but it also raises compute time, which is why moderate sizes work well for routine optimization [29]. The algorithm employs a beam size of 5 (gives a strong quality/latency trade-off for routine use) with a length penalty factor of 1.0 (defaults), though beam size 10 and length penalty 1.2 are shown in examples to balance sequence quality with computational efficiency. These values were chosen because they provided stable performance in testing while keeping runtime low [30]. Full decoding configurations could be found in the open-source implementation of ColiFormer. For *E. coli* optimization, we enforce GC bounds of 50-56%, providing a ± 2% tolerance around the optimal value to reflect natural sequence variability.

### 2.7. Statistical Analysis

For our metric, we report the mean ± standard deviation (SD). Comparisons between the original sequences and each optimization method were assessed with paired two-sided t-tests across matched genes. We also report p values (the probability of observing t at least this extreme if the true mean difference were zero), where t is a number that measures how far the observed mean difference is from zero in standard error units. For NCRF, the sample size is n = 61 due to its sequence length constraints; all other methods use n = 80. A significance threshold of α = 0.05 was used.

### 2.8. Code Availability

The open-source implementation of ColiFormer is publicly available at: https://github.com/SAKETH11111/ColiFormer. To facilitate broader adoption, we also offer a user-friendly interface for optimizing *E. coli* sequences, available at: https://huggingface.co/spaces/saketh11/ColiFormer.

## 3. Results

### 3.1. Optimizing Native E. Coli Genes

To determine ColiFormer’s ability to optimize both native *E. coli* sequences and established recombinant targets, we benchmarked it on 37,053 native *E. coli* genes alongside 80 industry-relevant genes derived from past recombinant protein expression studies. In addition, to compare ColiFormer’s performance to current approaches, we evaluated it against established methods, including IDT, Extended Random Choice (ERC), NCRF, CodonGPT, and CodonTransformer.

After benchmarking on the native *E. coli* genes, ColiFormer demonstrated a significant improvement in the CAI, which measures how closely a gene’s codon usage matches that of highly expressed genes in a given organism, making it a strong predictor of gene expression levels in vivo [31]. The baseline CAI of 0.689 represents the average across the 37,053 non-high-expression *E. coli* genes used for evaluation. The average CAI increased from 0.689 in the original sequences to 0.904 after optimization, representing an approximate 31% improvement. This enhancement strongly suggests greater codon usage alignment with highly expressed genes, indicating likely improvements in recombinant protein yield and cellular expression efficiency. The CAI increase is shown in Figure 3.

**Figure 3.**
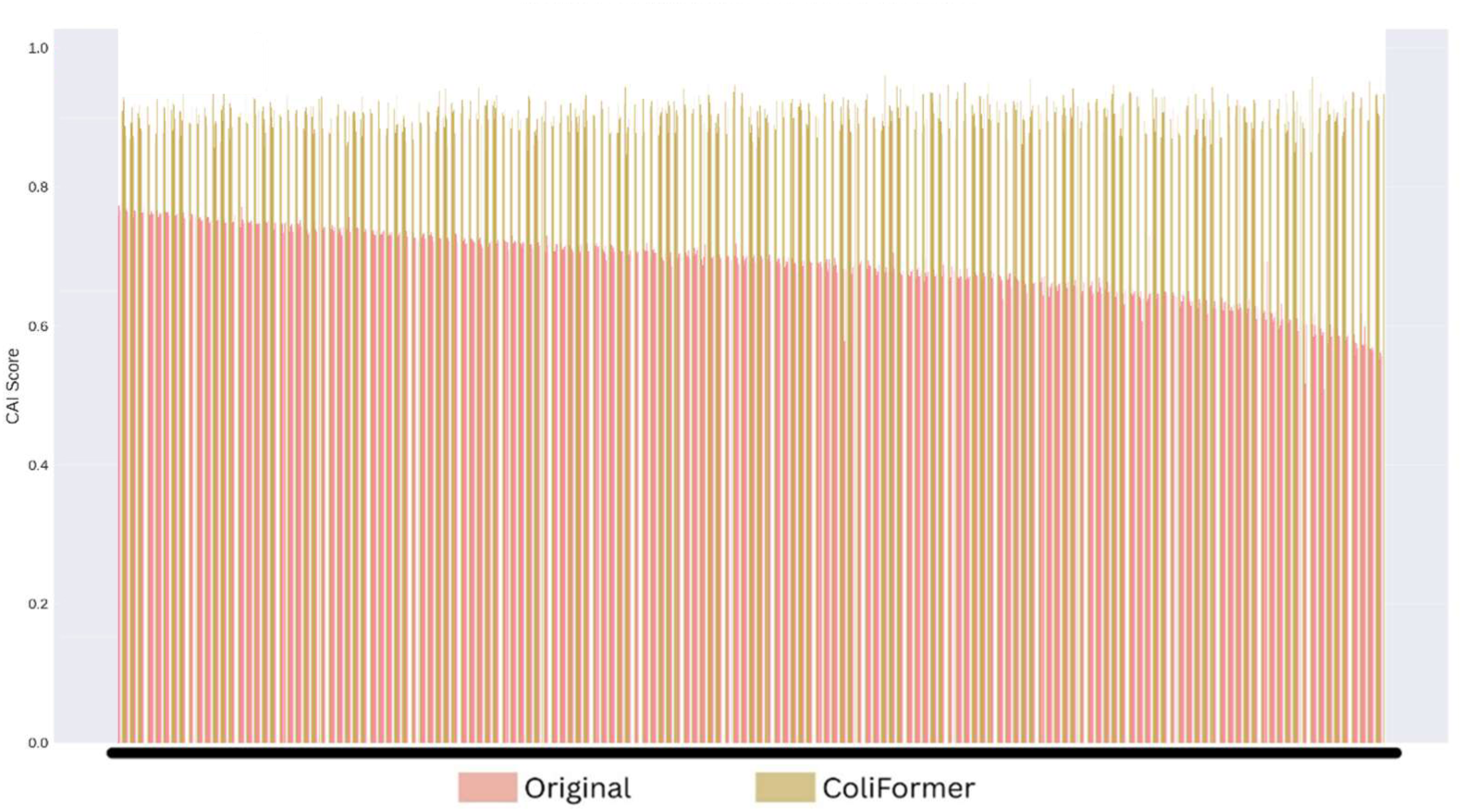
ColiFormer increased the CAI compared to the original native *E. coli* sequences (n=37,053).

Regarding the other key parameters affecting expression quality [31, 32], ColiFormer maintained the GC content within the optimal range, increasing it modestly from 51.24% to 55.96%. The increase to 55.96% remains within the biologically preferred 50-56% range for *E. coli*, avoiding extremes that affect expression efficiency. Importantly, the model reduced the incidence of negative cis-regulatory elements from 0.162 to 0.068. The tAI also increased from 0.303 to 0.373, further indicating stable protein expression. All individual values for each sequence before and after optimization across these key parameters are provided in Supplemental File 4 for further analysis.

### 3.2. Optimizing Established Recombinant Targets

To contextualize ColiFormer’s performance in optimizing established recombinant targets, IDT, CodonTransformer, NCRF, and ERC were used to optimize the benchmark set of the 80 industry-relevant genes. Those 80 genes have an average CAI of 0.58 ± 0.081. ColiFormer’s optimization yielded a CAI of 0.901 ± 0.0278 (p < 1×10⁻¹⁶), satisfying a 55.3% improvement compared to the original. ERC showed a modest improvement to 0.645 ± 0.083 (+11% over original); IDT optimization achieved an average CAI of 0.754 ± 0.033 (+30%); CodonTransformer reached 0.871 ± 0.029 (p < 1×10⁻¹⁶) (+50%); NCRF optimization yielded a CAI of 0.9634 (p < 1×10⁻¹⁶) (+66%). Although NCRF produced the highest average CAI of 0.9634 ± 0.0095, its success rate was only 76.3%. The 76.3% success rate reflects the model’s inability to process sequences longer than ∼800 amino acids due to input token limitations, with failures concentrated in 19 large proteins averaging ∼830 amino acids. The mean CAI for all approaches is shown in Figure 4.

**Figure 4.**
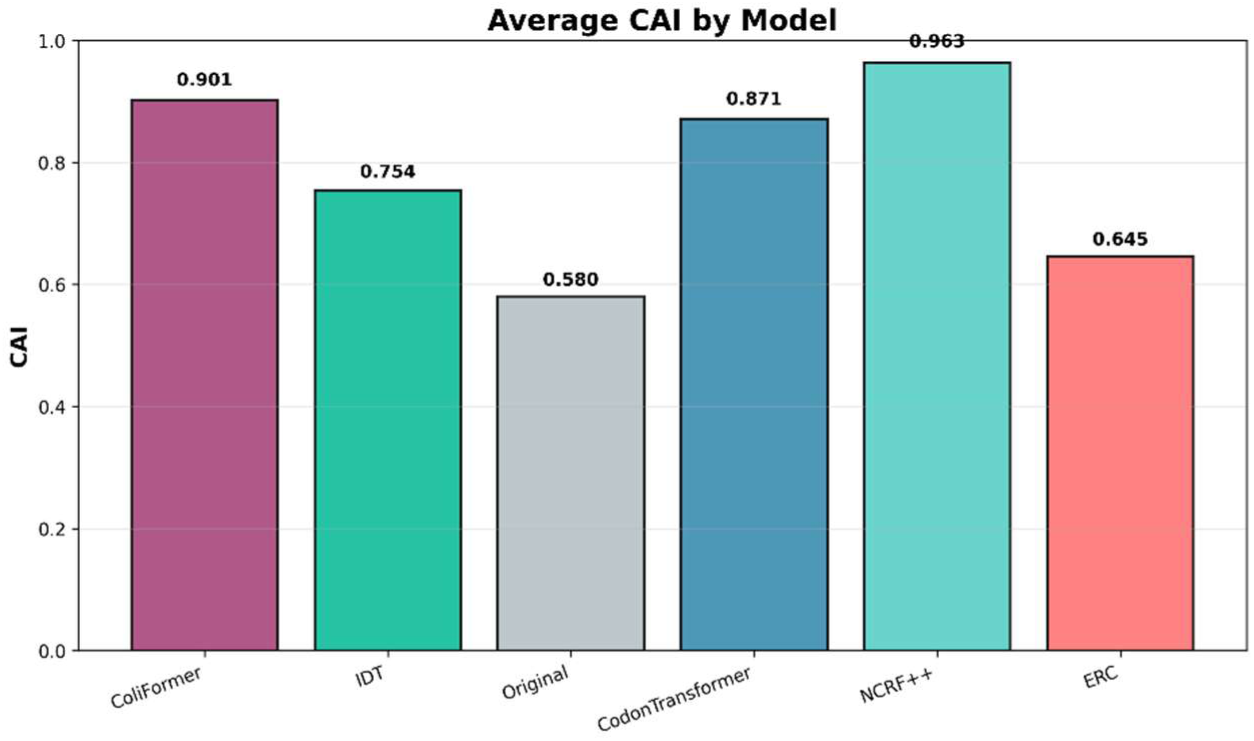
ColiFormer significantly increased the CAI compared to the IDT, Codon Transformer, and ERC techniques on the 80 industry-relevant genes (n=80).

Across the models, the optimal GC content range is generally accepted between 45–65%, with deviations outside this range negatively impacting molecular processes and gene performance [33, 34]. Specifically, the *E. coli* genome shows an optimal GC content between 52% and 56% [15, 35, 36]. ColiFormer maintains an average GC content of 56.38% ± 7.56% (p = 9.7e−10), very close to the optimal value, well within this accepted range, along with the other models. The mean GC Content for all approaches is shown in Figure 5.

**Figure 5.**
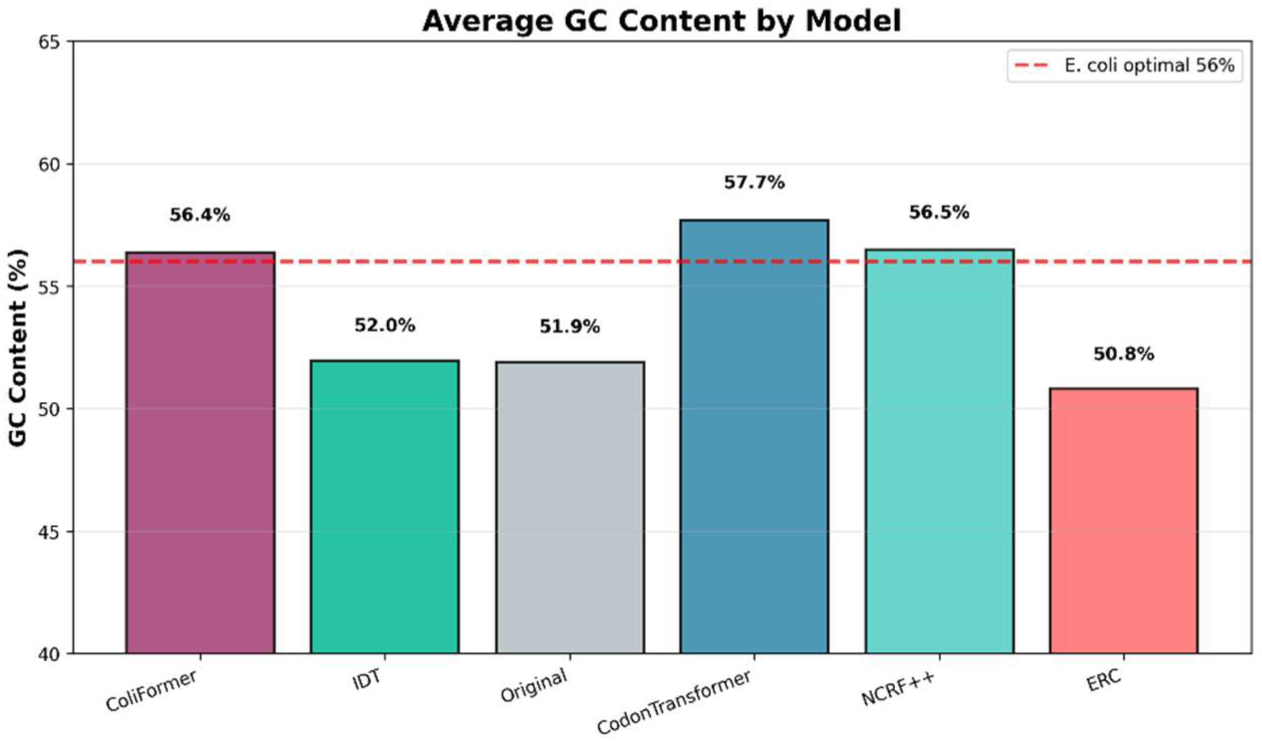
ColiFormer and the other codon optimization techniques maintained the GC content in the accepted range, with ColiFormer being closer to the *E. coli* optimal range than NCRF (56.5% ± 9.18%) (p = 5.6e−08) and CodonTransformer (57.7 ± 7.76) (p = 5.3e−14) on the 80 industry-relevant genes (n=80).

The tRNA Adaptation Index measures how well a gene’s codon usage matches the abundance and efficiency of the cellular tRNA pool, with higher tAI values indicating better translational efficiency [31]; ColiFormer achieved a high average tAI of 0.3628 ± 0.032 (p < 1×10⁻¹⁶) (27% over the original), notably higher than the unexpected tAI of NCRF and CodonTransformer. The mean tAI for all approaches is shown in Figure 6.

**Figure 6.**
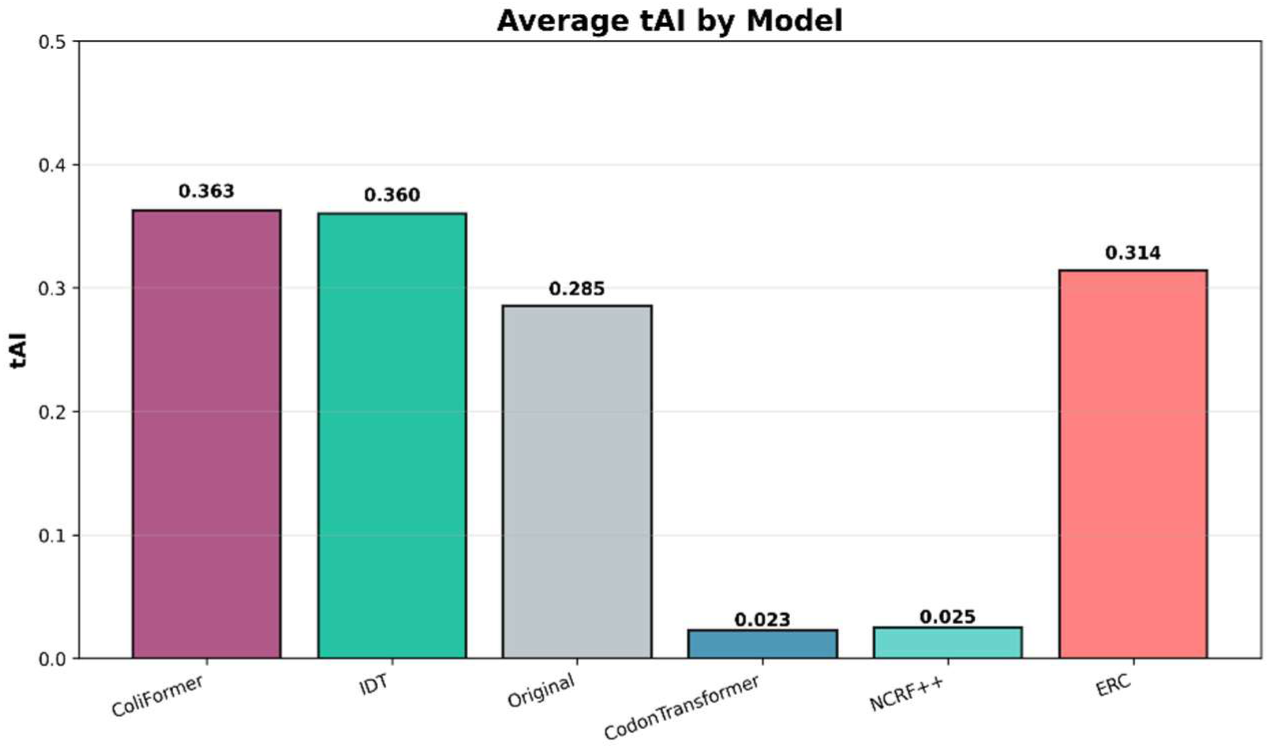
ColiFormer significantly increased the tAI compared to the IDT (0.36 ± 0.029) (p < 1×10⁻¹⁶), CodonTransformer (0.0228 ± 0.0012), ERC (0.314 ± 0.0296), and NCRF (0.0249 ± 0.0014) techniques on the 80 industry-relevant genes (n=80).

RNA stability was assessed by calculating the change in folding energy (Δ5’ MFE) over the initial 50 nucleotides. This metric impacts ribosome binding and translation initiation efficiency, where lower values correspond to enhanced translational efficiency [37]. ColiFormer achieved a Δ5’ MFE of 0.22 kcal/mol, representing the closest to the native folding energy of the original sequences, indicating the highest preservation of native RNA folding structure. This performance exceeded that of Codon Transformer (0.31 kcal/mol) and ERC (0.70 kcal/mol). In contrast, NCRF exhibited the lowest mean Δ5’ MFE (−2.49 kcal/mol), suggesting the strongest structural stabilization, while IDT also lowered the 5’ MFE (−0.40 kcal/mol) relative to the original sequence baseline (0.00 kcal/mol). The mean change in folding energy (Δ5′ MFE) over the first 50 nucleotides for all approaches is shown in Figure 7.

**Figure 7.**
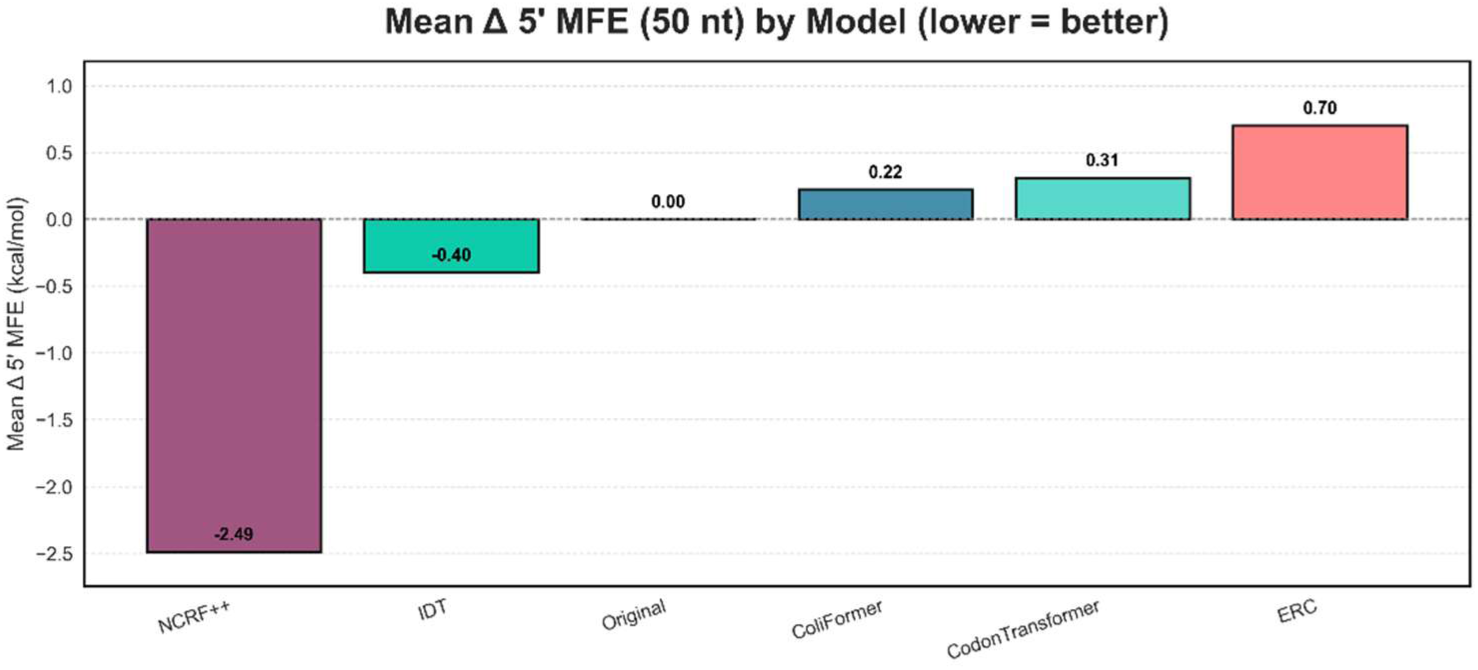
NCRF showed the greatest stabilization, while ColiFormer showed the best preservation of the native RNA structure, compared to CodonTransformer and ERC techniques on the 80 industry-relevant genes (n=80).

Negative cis-regulatory elements act as inhibitory sequences that can severely repress gene expression, and their decrease is beneficial for effective transcription and translation [38]. It includes known inhibitory motifs such as RBS occlusion sequences and premature poly-U tracts. ColiFormer demonstrated the lowest average negative cis elements of 0.0625, substantially lower than the other techniques, with values ranging from 3.06 to 5.26. This highlights ColiFormer’s strength in balancing multiple critical factors. The mean negative cis-regulatory elements for all approaches are shown in Figure 8.

**Figure 8.**
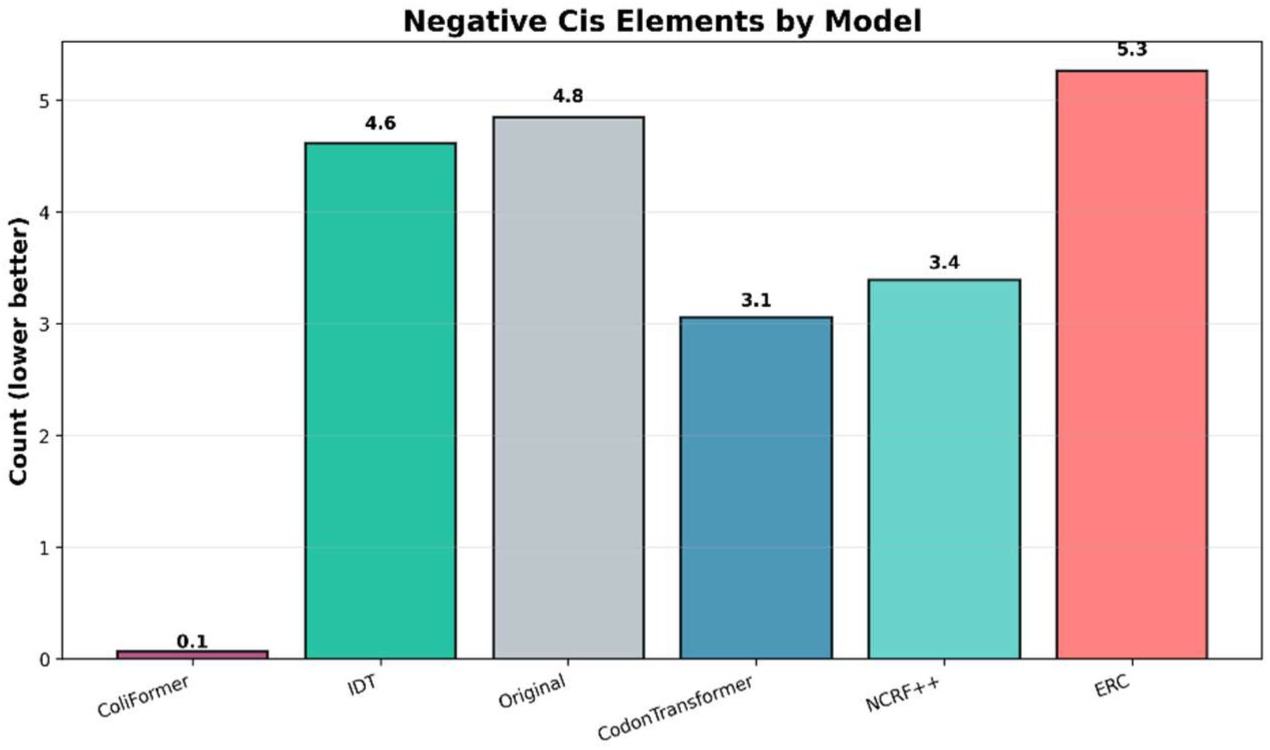
ColiFormer had the lowest negative cis-regulatory elements compared to the IDT, Codon Transformer, ERC, and NCRF techniques on the 80 industry-relevant genes (n=80). This demonstrates ColiFormer’s multi-parameter optimization capability.

The run time was calculated for ColiFormer and the ERC, CodonTransformer, and NCRF techniques on the benchmark set of 80 sequences. However, the average runtime for IDT was not calculated because it is a web-based tool rather than a standalone software or code that can be timed directly. The scores are displayed in Table 1.

**Table 1.**
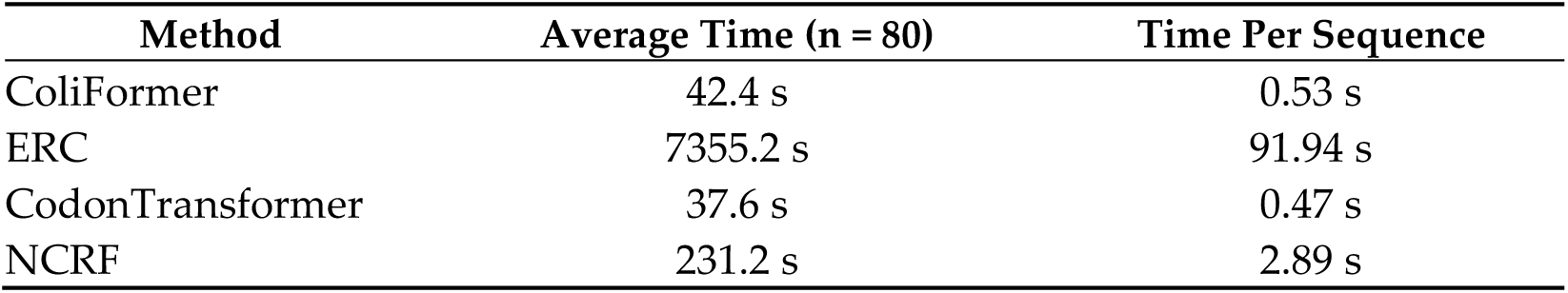
ColiFormer’s fast run time enables practical use on standard desktop GPUs, making it suitable for real-time design workflows, compared to ERC and NCRF. CodonTransformer shows similar runtime performance due to shared architectural components, while ColiFormer maintains comparable speed despite additional constraint handling.

| Method | Average Time (n = 80) | Time Per Sequence |
| --- | --- | --- |
| ColiFormer | 42.4 s | 0.53 s |
| ERC | 7355.2 s | 91.94 s |
| CodonTransformer | 37.6 s | 0.47 s |
| NCRF | 231.2 s | 2.89 s |

In summary, our comprehensive experiments demonstrated that ColiFormer consistently achieves superior performance compared to existing codon optimization methods. This advantage is reflected in improved benchmark metrics, including CAI, tAI, and GC content control, motivating the discussion of ColiFormer’s innovations and impact in the following section. All individual readings and optimization results for each technique on the benchmark set of 80 sequences are provided separately as follows: ColiFormer in Supplemental File 5, CodonTransformer in Supplemental File 6, NCRF in Supplemental File 7, ERC in Supplemental File 8, and IDT in Supplemental File 9.

## 4. Discussion

This study introduced ColiFormer, a transformer-based codon optimization model specifically fine-tuned for E. coli, that integrates multi-objective optimization to enhance recombinant gene expression. Across all benchmarks, ColiFormer outperformed existing tools, achieving higher CAI, improved tAI, and substantially fewer negative cis-regulatory elements, while maintaining biologically appropriate GC ranges. These results highlight ColiFormer’s ability to improve predicted expression without imposing undue cellular burden, demonstrating that effective codon optimization requires balancing expression efficiency with biological compatibility.

A key innovation of ColiFormer is its augmented-Lagrangian approach for precise GC-content control, which directly incorporates GC constraints into the model’s objective function rather than relying on post-hoc filtering. This integrated strategy enables dynamic adjustment of codon probabilities based on projected GC deviation, providing tighter and more biologically grounded control, a feature that is largely absent in existing models. In addition to its training-time advantages, the constrained beam search used during sequence generation improves reliability compared with standard beam search by preventing GC drift under strong optimization pressure.

A center biological insight of this work is that codon optimization must be balanced with cellular compatibility to preserve natural translation dynamics. Over-optimization toward highly abundant codons is known to disrupt natural ribosomal pausing and trigger proteotoxic stress, all of which can impair co-translational folding and reduce soluble protein yield. Prior studies show that harmonization of rare codons preserves essential pauses required for domain-wise folding and mitigates cellular stress responses. By explicitly modeling both optimization and harmonization, ColiFormer avoids the classical trade-off where high CAI sequences exhibit misfolding or metabolic burden. This supports emerging evidence that biologically informed codon design is essential for maintaining translational homeostasis and cell viability.

An additional strength of ColiFormer lies in its augmented Lagrangian GC-control strategy, which provides practical and reproducible advantages over conventional codon optimization tools. Most existing approaches either ignore GC content entirely or apply simple post-hoc filters that do not interact with the optimization process. In contrast, ColiFormer integrates GC constraints directly into training and sequence generation, allowing codon choices to be dynamically adjusted based on projected GC deviation. This results in tighter, more biologically relevant control compared to rule-based or heuristic methods, which often drift toward extreme GC content values when aggressively optimizing CAI.

As outlined earlier in the Introduction, we initially intended to benchmark ColiFormer against the RL (Reinforcement Learning)-optimized variant of CodonGPT, which reportedly achieves CAI values between 79–84% [19]. However, the RL-trained CodonGPT model has not been made publicly available, precluding direct benchmarking. Despite this, ColiFormer achieves CAI scores higher than those reported for CodonGPT across our test set.

From a computational perspective, ColiFormer demonstrates a low computational complexity, optimizing sequences in approximately 0.5 seconds each. This remarkable processing speed is a major advantage over many existing methods that require considerably longer runtimes, enabling highly scalable workflows suitable for industrial and research applications requiring high-throughput gene design

Our evaluation framework incorporates a comprehensive set of in silico metrics. However, these metrics do not fully capture all factors influencing gene expression. Recent research highlights the potential to further enhance translation dynamics modeling by integrating metrics like elongation rate and tRNA adaptation, which could provide more nuanced insights into translational efficiency and protein folding [39]. ColiFormer’s architecture and output are well-suited for such extensions in future work. Because the augmented-Lagrangian objective treats each constraint as an independent, differentiable term, it can readily incorporate additional sequence features. Likewise, the constrained decoding procedure can be extended to enforce any property, such as maintaining desired mRNA secondary structure or controlling the usage of rare codons, enabling broader constraint-aware sequence design in future versions of ColiFormer.

Looking ahead, ColiFormer can be extended in several impactful ways. Incorporating models of elongation rates and codon-pair context would capture translation dynamics more accurately, while integrating structural protein modeling could link codon usage to folding stability and function. Likewise, experimental validation of optimized sequences will be essential to confirm computational predictions and guide further refinement in future studies.

Regarding codon usage metrics, the CAI used in this study is functionally equivalent to the Codon Similarity Index (CSI), which applies CAI genome-wide to avoid bias from selecting specific reference genes. This ensures optimization reflects overall *E. coli* codon usage rather than overemphasizing highly expressed genes, which can sometimes disrupt folding fidelity. However, CAI/CSI have limitations: they assume that codon patterns in highly expressed genes always predict optimal translation, which overlook codon-pair effects and mRNA structural constraints.

ColiFormer’s multi-objective design helps mitigate these issues and remains extensible for future integration of more mechanistic translation metrics. Although ColiFormer is currently trained exclusively on *E. coli* genes, its methodology is broadly adaptable for other host organisms, including yeast and mammalian cells. Transfer learning approaches can leverage pre-trained weights and fine-tune the model using organism-specific data, accelerating adaptation.

Finally, to ensure transparency and reproducibility, we emphasize the importance of making the ColiFormer model and its codebase publicly available. Open access to these resources allows independent verification of results and supports broader adoption of the methods introduced in this study. By enabling the community to retrain and extend ColiFormer, we help establish stronger standards for codon optimization research and ensure long-term usability of the tool.

It is important to note that experimental results of ColiFormer-optimized sequences are not yet included. While this study contributes primarily to computational methods, future work will involve experimental validation of ColiFormer-optimized sequences in *E. coli* expression systems to confirm the predicted improvements in vivo and assess potential regulatory effects.

## Supplementary Materials

The following supporting information is available online:

Supplemental File 1: Clustered *E. coli* gene sequences (CD-HIT-EST output).

Supplemental File 2: Final training dataset gene sequences.

Supplemental File 3: Benchmark set of 80 recombinant genes.

Supplemental File 4: Individual optimization parameter values for native *E. coli* genes.

Supplemental File 5: ColiFormer optimization results on benchmark sequences.

Supplemental File 6: CodonTransformer optimization results.

Supplemental File 7: NCRF optimization results.

Supplemental File 8: ERC optimization results. Supplemental File 9: IDT optimization results.

## Author Contributions

S.B. and O.E. were involved in determining the datasets and methods of comparison for codon optimization tools under the supervision of A.E. and G.L.; O.E., S.B., F.C., I.F., and Y.S. were involved in the writing and reviewing of the manuscript and in the interpretation of the results. O.E. and A.E. were involved in administration. The first two authors made equal contributions to this work. All authors edited, revised, read, and approved the published version of the manuscript.

## Institutional Review Board Statement

Not applicable

## Informed Consent Statement

Not applicable

## Data Availability Statement

The *E. coli* gene sequences and the 80-benchmark set used in this study were obtained from the National Center for Biotechnology Information (NCBI) GenBank database [20]. The ColiFormer model is built upon the CodonTransformer architecture [18]. The curated datasets generated and/or analyzed during the current study, including training and benchmark sets, are available in the publicly accessible repository GitHub at https://github.com/SAKETH11111/ecoli. Supplementary files containing additional data related to benchmark genes and optimization results are provided with the article. All code and models used for ColiFormer development are also publicly available through the repository to enable reproducibility.

## Acknowledgments

Not applicable

## Abbreviations

The following abbreviations are used in this manuscript:

E. coli: Escherichia coli
CAI: Codon Adaptation Index
tAI: tRNA Adaptation Index
GC: Guanine-Cytosine (content)
NCBI: National Center for Biotechnology Information
MLM: Masked Language Modeling
IDT: Integrated DNA Technologies
ERC: Extended Random Choice
NCRF: Neural Codon Random Forest
STREAM: Shared Token Representation and Encoding with Aligned Multi-masking
UNK: Unknown token (used in sequence tokenization)
tRNA: Transfer Ribonucleic Acid
VRAM: Video Random Access Memory (GPU memory)
RL: Reinforcement Learning

